# STRUCTURE-ACTIVITY STUDIES ON ANALOGUES OF 4-METHYLGUAIACOL, A CATTLE ANAL ODOUR CONSTITUENT REPELLENT TO THE BROWN EAR TICK *(Rhipicephalus appendiculatus)*

**DOI:** 10.1101/460725

**Authors:** Margaret W. Kariuki, Ahmed Hassanali, Margaret M. Ng’ang’a

**Affiliations:** Department of Chemistry, School of Pure and Applied Sciences, Kenyatta University, P.O. Box 43844-00100, Nairobi.

**Keywords:** Key words, *Rhipicephalus appendiculatus*, structural analogues, 4-methylguaiacol, 4-propylguaiacol, repellence.

## Abstract

Previously, 4-methylguaiacol, a major constituent of cattle anal odour, was found to have a high repellence on *Rhipicephalus appendiculatus.* In the present study, 10 structural analogues of the phenol were tested for repellence against *R. appendiculatus* in order to assess the effects of (i) absence or presence of the 4-alkyl group of varying length, (ii) inclusion of a double bond in the 4-alkyl chain, (iii) linking the two phenolic oxygen in a methylenedioxy bridge, (iv) replacement of the OCH_3_ with CH_3_ and inclusion of another CH_3_ at position 6, and (v) presence of an additional OCH_3_ group at position 6. The analogues comprised of 2-methoxyphenol (guaiacol), 4-ethyl-2- methoxyphenol, 4-propyl-2-methoxyphenol, 4-allyl-2-methoxyphenol (eugenol), 3,4-methylenedioxytoluene, 2,4- dimethylphenol, 4-ethyl-2-methylphenol, 2,4,6-trimethylphenol, 4-propyl-2,6-dimethoxy-phenol and 4-allyl-2,6- dimethoxyphenol, which were compared at different doses in a two-choice climbing assay set up. Each analogue showed either increased or reduced repellence compared with 4-methylguaiacol. The structural feature that was associated with the highest repellence was 4-propyl moiety in the guaiacol unit (RD_75_ = 0.031 for 4-propyl-2- methoxyphenol; that of 4-methylguaiacol = 0.564). Effects of blending selected analogues with high repellence were also compared. However, none of the blends showed incremental increase in repellence compared with that of 4- propyl-2-methoxyphenol. We are currently evaluating the effects of controlled release of the compound at different sites on cattle on the behavior and success of *R. appendiculatus* to locate their predilection feeding site.

## Introduction

East Coast fever (ECF), caused by *Theileria parva parva* (Theiler, 1904), and transmitted by the brown ear tick, *R. appendiculatus* (Neumann, 1901), is one of the major constraints in the development of the livestock industry in eastern and southern Africa (Olwoch *et al.,* 2008; Fry *et al.,* 2016). Of the estimated 12.7 million heads of cattle (both indigenous and exotic), 76% are at risk to ECF (Lawrence *et al.,* 1996). The disease is associated with up to 10% mortality in zebu calves in ECF endemic areas and can cause up to 100% mortality in susceptible exotic and indigenous breeds (Mbogo, *et* al.,1995; Lawrence *et al.,* 1996; Gachohi *et al.,* 2012).

On host behavior studies of adult *R. appendiculatus* shows preference to feed mainly inside and around the ear of their hosts (Wanzala *et al.,* 2004). A repellent blend from the anal region and an attractive blend at the ear have been shown to play natural “push” and “pull” roles, respectively, to guide these ticks to the cattle ears (Wanzala *et al.,* 2004). Interestingly, application at the cattle ears of two tick-repellent essential oils from ethno-plants growing in western Kenya were found to confuse the ticks, most of which dropped off the host animals (Wanzala *et al.,* 2018).

In a preliminary study, crude odour collected from the cattle anal region was placed at the ears of several Friesian steers. Most of the brown ear ticks released at different sites of the animals also failed to locate their predilection sites and dropped off to the ground. Recently, the major constituents of the odour were characterized and assayed in climbing assays (Kariuki *et al.,* 2018). The most active compound was found to be 4-methylguaiacol. Interestingly, in a previous structure-activity study with different analogues of tsetse repellent guaiacol (Torr *et al.,* 1996), 4- methylguiacol was also found to be more repellent to savannah tsetse than the parent compound and other analogues (Saini and Hassanali, 2007). The objective of the present study was to determine if different structural variants of 4- methylguaiacol affected their repellence to *R. appendiculatus.* In addition, effects of blending selected more repellent analogues were also studied to see if there were any additive or synergistic effects between some of the compounds.

## Materials and methods

### Ticks

The ticks (brown ear tick, *R. appendiculatus* Neumann, 1901) were obtained from the colonies at the International Livestock Research Institute (ILRI),Nairobi, Kenya. Rearing conditions and management of ticks were as described previously (Irvin & Brocklesby, 1970). All the experiments were conducted using newly emerged adult ticks of mixed sexes

### Tested compounds

The choice *of* test compounds (**1-11)** was based on structural modification as per the following criteria:

i. absence or presence of the 4-alkyl group of varying length (compounds 1-4);
ii. inclusion of a double bond in the 4-alkyl chain (compound 5);
iii. linking the two phenolic oxygen in a methylenedioxy bridge (compound 6);
iv. replacement of the OCH_3_ with CH_3_ and inclusion of another CH_3_ at position 6 (compounds 7-9);
v. and presence of an additional OCH_3_ group at position 6 (compounds 10 and 11).

The test compounds shown in figure 1 were of analytical grade and obtained from Sigma-Aldrich Company (Germany). Each compound was tested at different doses against untreated control.

**Figure.1.**
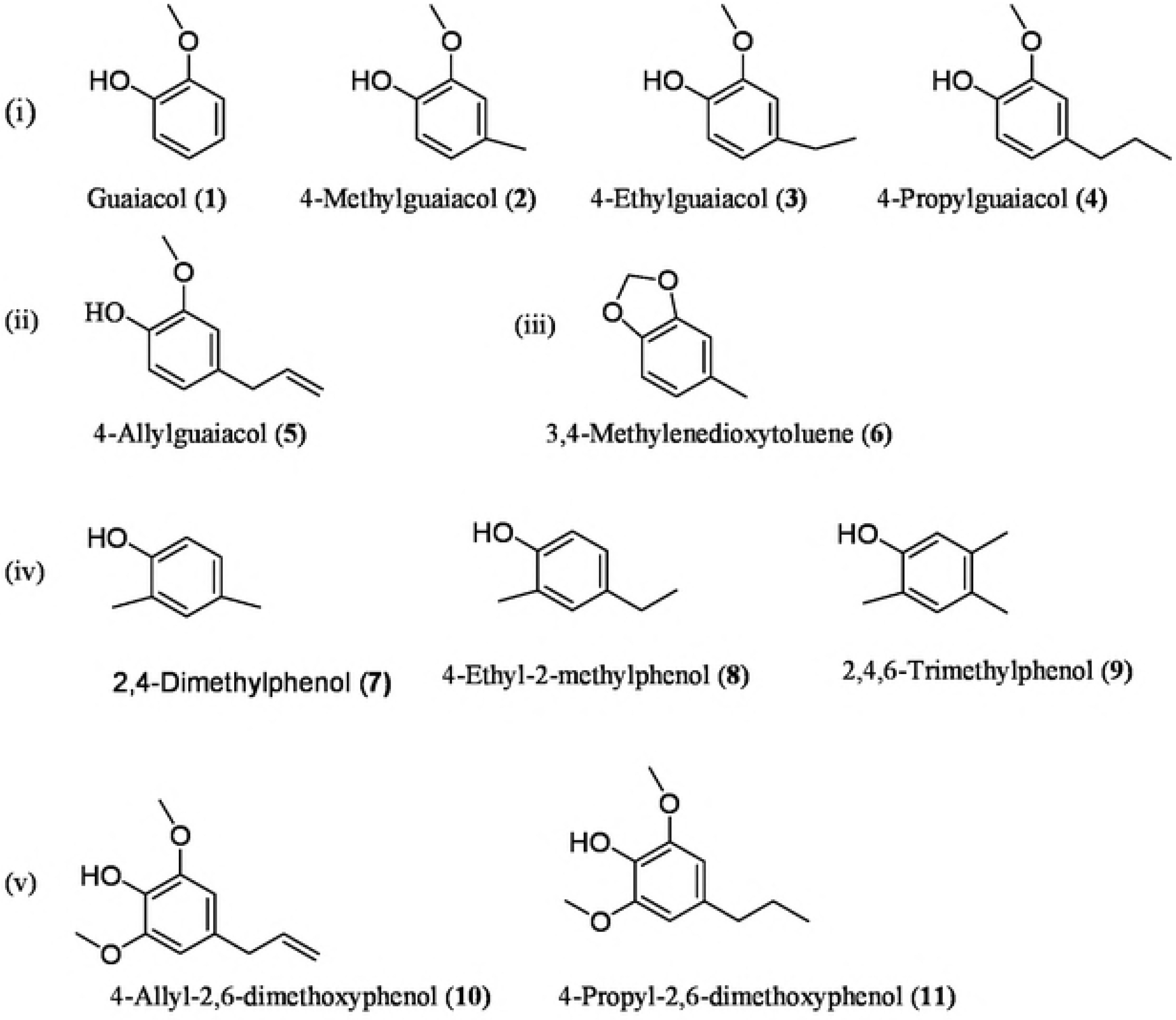
Structures of 4-methylguaiacol and selected analogues evaluated.

A blend containing the four most potent compounds (4-propyl guaiacol (4), 4-allyl-2,6-dimethoxyphenol, 2,4- dimethylphenol and 4-propyl-2,6-dimethoxyphenol) was constituted and also tested against the untreated control, followed by subtractive bioassays of the same with each of the four compounds missing in the blend.

### Dual-Choice Repellence Assays

A dual-choice tick repellence climbing assay (Wanzala *et al.,* 2004) that exploits the behaviour of *R. appendiculatus* to climb up grass stems to await potential hosts passing by was used (Chiera, 1985). The repeUence of 4- methyl-guaiacol and the 10 analogues against *R. appendiculatus* were compared. In addition, blends of four most active compounds (in equal proportions) and four blends with each of the four components missing were also tested. Each compound and blend were diluted serially with dichloromethane (analytical grade) to provide 10 to 0.01 mg/ml solutions. An aliquot of 100μl of each dose was applied to a filter paper strip on the glass tube, with an equivalent volume of dichloromethane added to the control filter paper strip. The set up was allowed to equilibrate for 30 min before five adult brown ear ticks of mixed age and sex were released at the base of the climbing assay set-up. Observations were made over a 1-hour period, and the numbers of ticks above the filter paper strip on the control glass tube (Nc) and on the glass tube with test materials (Nt) were recorded at 15, 30, 45, and 60 min. Six replicates for each dose were carried out, each time with fresh, nave adult ticks. Initial comparison of the responses of ticks in the set-up with and without residual dichloromethane on both sides showed no bias for either side and no effects of the residual solvent on the adult ticks. The repellency of each dose was calculated using the formula: number of ticks in the control arm - number of ticks in the treated arm/total responding ticks) × 100. Dose-response % repellency data from the replicated experiments were subjected to probit analysis:

Probit [P(Conc.1]=β_0_ +x β_j_ + î, where β_0_ = coefficient of the model representing y-intercept,

β_1_ = coefficient of the model representing conc. 1,

conc. 1 =log_10_ (conc.),

î =error term in the data set of the predictor (regressor) variable (x) and

p = repellency probability

### Data analysis

The repellency data obtained at different concentrations for each compound and blend were subjected to analysis of variance (ANOVA) for a completely randomized design. Dose-response relationship was determined using probit analysis and repellent dose at RD_75_ values obtained from the regression model treatment means were separated using Student-Newman-Keuls (SNK) at p ≤ 0.05 significance level.

## Results

The responses of *R. appendiculatus* to each compound tested in the two-choice climbing assay at different doses are summarized in Table 1.

**Table 1.**
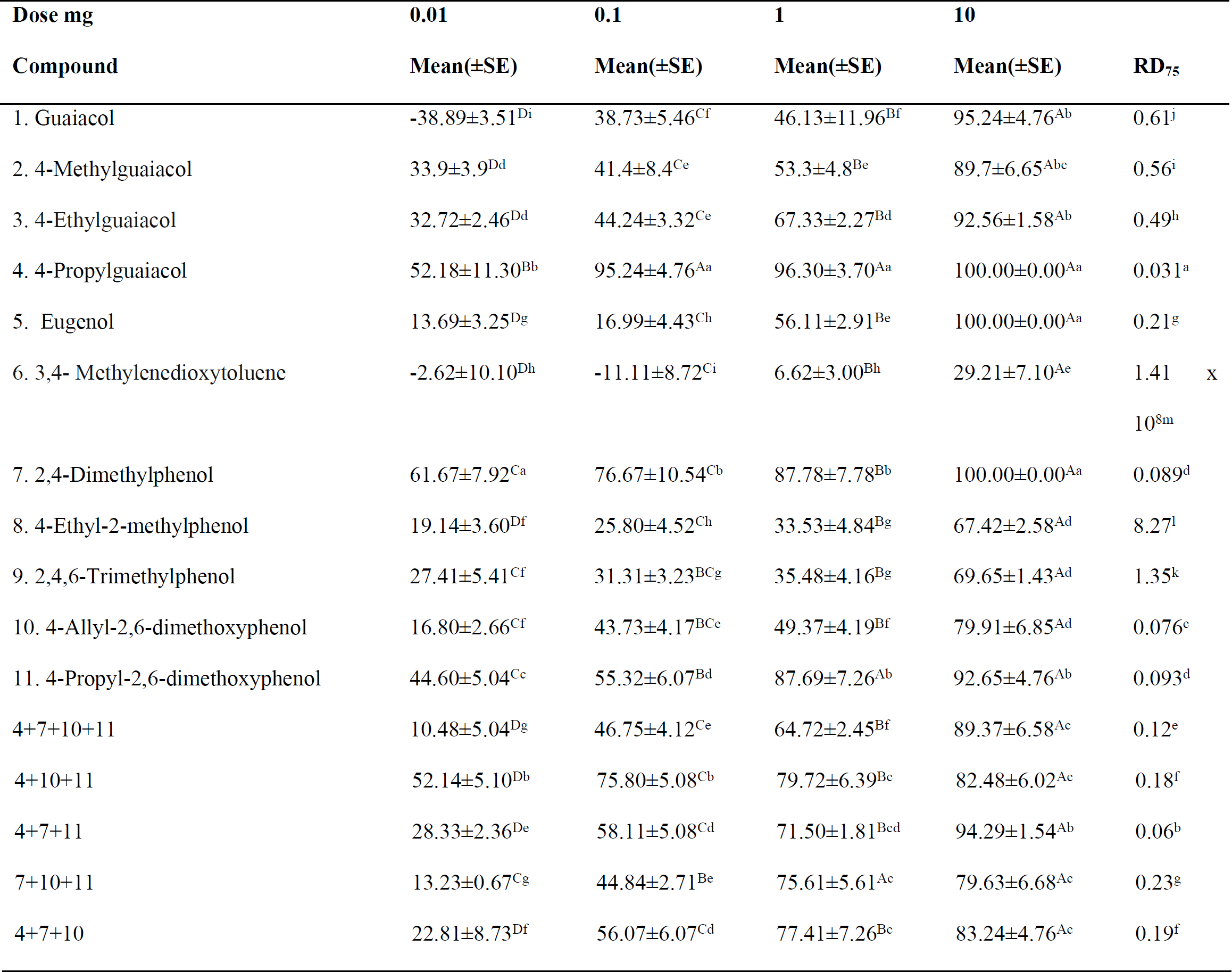
THE REPELLENT EFFECTS OF THE ANALOGUES OF 4-METHYLGUAIACOL AND SELECTED BLENDSBASED ON THE 4 MOST POTENT ANALOGUES

Mean (±SE) with the same lowercase letter in each column and uppercase letters in each row are not significantly different at α=0.05 (Student-Newman-Keuls test), respectively

4-Propylguaiacol exhibited the highest repellence (RD_75_ = 0.031) to the brown ear tick among all the compounds tested in the study. Other analogues that showed relatively high repellence include 4-allyl-2,6-dimethoxyphenol (RD _75_= 0.076), 2,4-dimethyl phenol (RD_75_ = 0.089), and 4-propyl-2,6-dimethoxyphenol (RD_75_ = 0.093). Of the blends tested, combination of 4-propylguaiacol, 2,4-dimethylphenol and 4-propyl-2,6-dimethoxyphenol showed the highest repellence (RD_75_ = 0.06), although lower than that of 4-propylguaiacol. Other blends showed levels of repellence that were lower than those of the individual compounds.

## Discussion

In this study, we compared the effects of different structural modifications of 4-methylguaiacol on their repellence to the brown ear tick, *R. appendiculatus.* Our overall objective was to identify more repellent candidates that can be deployed on-host at reduced release rates to effectively confuse the ticks and substantially reduce the numbers that successfully locate their predilection feeding sites. In our first set of comparison with the 4-alkyl moiety of the guaiacol unit, the results showed significant increases in the repellent effect with increasing length of the carbon chain (4-propylguaiacol > 4-ethylguaiacol > 4-methylguaiacol > guaiacol). 4-Propylguaiacol was found to be the most repellent analogue among all the compounds tested in the present study (with RD_75_ = 0.031 compared to RD_75_ = 0.56 of 4-methylguaiacol). However, replacement of the propyl chain with a double bond carrying allyl moiety in eugenol decreased the repellence (RD_75_ of eugenol = 0.21), showing that the flexibility of the hydrophobic carbon chain has a significant effect on its repellence to the tick. It will be interesting to see if further increases in the carbon chain of the 4-alkyl group (e.g. butyl and pentyl groups) have incremental effects on the repellence of the compounds.

Our study also revealed a number of interesting structural effects on levels of repellence. The absence of phenolic OH group in 3,4-methylenedioxytoluene led to a substantial drop (RD_75_ = 1.41 × 10^8^) in repellence (compared with that of 4-methylguaiacol) showing an important role it plays in repellence. On the other hand, replacement of the OCH3 of 4- methylguaiacol with CH_3_ showed significantly higher repellence of 2,4-dimethylphenol (RD_75_ = 0.089). However, inclusion of another CH_3_ at position 6 significantly reduced the repellence of 2,4,6-trimethylphenol (RD_75_ = 1.35). Interestingly, the presence of an additional OCH_3_ group at position 6 significantly raised the repellence of 4-allyl-2,6- dimethoxyphenol its repellence (RD_75_ = 0.076) relative to that (RD_75_ = 0.21) of 4-allyl-2-methoxyphenol (eugenol). On the other hand, its presence in 4-propyl-2,6-dimethoxyphenol reduced somewhat its repellence (RD_75_ = 0.093) relative to that of 4-propylguaiacol (RD_75_=0.031). These findings show variations in the effects of some substituents on the repellence of some analogues of 4-methylguaiacol to the brown ear tick, but also suggest other possible structural variants that would be interesting to evaluate.

In the present study, we also explored possible synergistic or additive effects of several (four or three) more repellent analogues (4-propylguaiacol, 2,4-dimethylphenol, 4-allyl-2,6-dimethoxyphenol and 4-propyl-2,6-dimethoxyphenol). However, none of the blends showed repellence comparable to that of 4-propylguaiacol. Repellence of 3-component blend without 4-allyl-2,6-dimethoxyphenol was closest (RD_75_ = 0.06) to that of 4-propylguaiacol. As expected, that of 3 components without 4-propylguaiacol was least repellent (RD_75_ = 0.23).

In summary, the present study on different structural variants of 4-methylguaiacol, a constituent of cattle anal odour repellent to the brown ear tick (Kariuki et al, 2018), identified 4-propylguaiacol as the most repellent analogue. We are currently evaluating the effect of microencapsulated controlled-release of this analogue at cattle ears on ticks placed at different location on the animal. The results of the study will be reported elsewhere.

### Ethics statement

The authors declare no conflicts of interest

## Acknowledgements

This work was supported by funds from MOHEST/ADB in collaboration with Kenyatta University and National Research Fund, ILRI and KARLO for provision of ticks and Chemistry department technicians, Kenyatta University for technical support.

## Ethical approval

All applicable international, national, and/or institutional guidelines for the care and use of animals were followed. This article does not contain any studies with human participants performed by any of the authors.

## REFERENCES

Chiera JW, Newson RM, Cunningham, MP. Cumulative effects of host resistance on Rhipicephalus appendiculatus Neumann (Acarina: Ixodidae) in the laboratory. Parasitol. 1985; 90: 401–409.

Fry LM, Schneider DA, Frevert CW, Nelson DD, Morrison WI, Knowles DP, East coast fever caused by theileria parva is characterized by macrophage activation associated with vasculitis and respiratory failure. PloS One. 2016; 11: e0156004.

Gachohi J, Skilton R, Hansen F, Ngumi P, Kitala P,. Epidemiology of East Coast fever (Theileria parva infection) in Kenya: past, present and the future. Parasit Vectors. 2012; 5: 194.

Irvin AD, Brocklesby DW. Rearing and maintaining Rhipicephalus appendicutatus in the laboratory. Inst Anim Tech J. 1970; 21: 106.

Kariuki MW, Ng’ang’a MM, Hassanali A. Characterisation of cattle anal odour constituents associated with the repellency of *Rhipicephalus appendiculatus.* Exp. Appl. Acarol. 2018; 76: 221–227.

Lawrence J.A, Musisi FL, Mfitilodze MW, Tjornehoj K, Whiteland AP, Kafuwa PT, Chamambala, KE. Integrated tick and tick-borne disease control trials in crossbred dairy cattle in Malawi. Trop. Anim. Health Prod. 1996; 28: 280–288.

Mbogo KS, Kariuki PD, McHardy, N. & Payne, R. (1995). Training Manual For Veterinary Staff on Immunization Against East Coast Fever Using the ECFiM System. Kenya Agricultural Research Institute and Overseas Development Administration of UK.

Olwoch JM, Reyers B, Engelbrecht FA, Erasmus BFN. Climate change and tick-borne disease, Theileriosis (East Coast Fever) in Sub-Saharan Africa. J. Arid Environ. 2008; 72: 108–120.

Saini RK, Hassanali A. A 4-alkyl-substituted analogue of guaiacol shows greaterrepellency to Savannah Tsetse (Glossina spp.). J. Chem. Ecol. 2007; 33: 985–995.

Torr SJ, Mangwiro TNC, Hall DR. Responses of *Glossina pallidipes (Diptera: Glossinidae) to synthetic repellents in the field*. Bull. Entomol. Res. 1996; 86: 609–616.

Wanzala W, Hassanali A, Mukabana WR, Takken W. The effect of essential oils of Tagetes minuta and Tithonia diversifolia on on-host behaviour of the brown ear *tickRhipicephalus appendiculatus*. Livestock Research for Rural Development. 2018; 30: 6.

Wanzala W, Sika NFK, Gule S, Hassanali A. Attractive and repellent host odours guide ticks to their respective feeding sites. Chemoecology. 2004; 14: 229–232.

